# Cul3 substrate adaptor SPOP targets Nup153 for degradation

**DOI:** 10.1101/2023.05.13.540659

**Authors:** Joseph Y. Ong, Jorge Z. Torres

## Abstract

SPOP is a Cul3 substrate adaptor responsible for degradation of many proteins related to cell growth and proliferation. Because mutation or misregulation of SPOP drives cancer progression, understanding the suite of SPOP substrates is important to understanding regulation of cell proliferation. Here, we identify Nup153, a component of the nuclear basket of the nuclear pore complex, as a novel substrate of SPOP. SPOP and Nup153 bind to each other and colocalize at the nuclear envelope and some nuclear foci in cells. The binding interaction between SPOP and Nup153 is complex and multivalent. Nup153 is ubiquitylated and degraded upon expression of SPOP^WT^ but not its substrate binding-deficient mutant SPOP^F102C^. Depletion of SPOP via RNAi leads to Nup153 stabilization. Upon loss of SPOP, the nuclear envelope localization of spindle assembly checkpoint protein Mad1, which is tethered to the nuclear envelope by Nup153, is stronger. Altogether, our results demonstrate SPOP regulates Nup153 levels and expands our understanding of the role of SPOP in protein and cellular homeostasis.

## Introduction

The Cullin family of proteins are E3 ubiquitin ligases responsible for about 20% of ubiquitin-mediated proteasomal degradation within a cell (Soucy et al., 2009). Cul3 assembles an active E3 ubiquitin ligase complex by simultaneously binding an E2 ubiquitin conjugating enzyme charged with the ubiquitin and a BTB-domain containing substrate adaptor which binds to the substrate to be ubiquitylated (Petroski and Deshaies, 2005). Prominent Cul3 substrate adaptors include Keap1, LZTR1, and SPOP. Keap1 binds to and targets the transcription factor Nrf2, a protein responsible for responses to oxidative conditions and cellular stress, for degradation (Cullinan et al., 2004; Furukawa and Xiong, 2005). LZTR1 binds to and targets the Ras family of GTPases (Steklov et al., 2018; Bigenzahn et al., 2018) and Ras-related GTPase RIT1 (Castel et al., 2019) for degradation.

SPOP is a Cul3 substrate adaptor with three main domains: a MATH domain that binds to substrates, a BTB domain that causes SPOP to dimerize and also mediates interaction with Cul3, and an oligomerization BACK domain through which SPOP dimers can chain to form higher-order structures (Marzahn et al., 2016). Hydrophobic residues within the MATH, such as Y87, F102, Y123, W131, and F133, mediate the interaction between SPOP and its substrates (Zhuang et al., 2009), and mutations of these residues generally disrupt the ability of SPOP to bind to its substrates (Geng et al., 2013a; Janouskova et al., 2017; Ostertag et al., 2019a). Because SPOP plays important roles in cell grow regulation, SPOP is commonly mutated or misregulated in cancers, suggesting an important role for SPOP as a tumor suppressor (Barbieri et al., 2012; Le Gallo et al., 2012; Hu et al., 2016; Wang et al., 2020b).

Whereas the principle substrate of Keap1 is Nrf2 and that of LZTR1 are Ras-related GTPases, SPOP regulates many proteins involved in cell growth and proliferation. For example, androgen receptor (AR)-mediated signaling regulates cell proliferation and differentiation transcriptional programs (Culig and Santer, 2014). SPOP mediates the degradation of AR (An et al., 2014; Geng et al., 2014), its co-activator SRC-3 (Geng et al., 2013b; Zhou et al., 2010), and TRIM24, an enhancer of AR-mediated gene activation (Groner et al., 2016), and SPOP mutations derived from prostate cancers disrupt SPOP’s ability to regulate AR-signaling. Similarly, SPOP targets estrogen-receptor for degradation in endometrial cancers (Zhang et al., 2015). In development, SPOP regulates Hedgehog signaling by degrading transcription factors Gli2 and Gli3 (Chen et al., 2009) and consequently influences morphogenesis in both *Drosophila* and vertebrates (Cai and Liu, 2016, 2017; Schwend et al., 2013; Seong and Ishii, 2013; Kent et al., 2006). Correspondingly, SPOP misregulation is associated with dysfunction of Hedgehog signaling in a variety of cancer types (Li et al., 2014; Zhi et al., 2016; Jin et al., 2019b; Zeng et al., 2014; Li et al., 2019; Burleson et al., 2022). In some cancer types, the cancer stem cell-associated transcription factors c-Myc (Luo et al., 2018; Geng et al., 2017) and Nanog (Wang et al., 2019; Zhang et al., 2019) are also degradation targets of SPOP. More broadly, wild-type SPOP promotes genomic stability, and loss of SPOP function can result in defects in DNA replication stress (Hjorth-Jensen et al., 2018), double-stranded break repair (Boysen et al., 2015), and the DNA-damage response (Jin et al., 2021). Altogether, given SPOP’s regulatory roles in cell growth, proliferation, and development, understanding the suite of SPOP substrates is crucial to understanding SPOP’s role in cell biology and its pathological roles when mutated or misregulated.

The human nuclear pore complex (NPC) is an approximately 120 MDa complex consisting of multiple copies of about 30 proteins termed nuclear pore complex proteins (Nups) (Beck and Hurt, 2016). As the main channel between the nucleoplasm and the cytoplasm, the NPC plays a crucial role in the trafficking of proteins and RNAs in and out of the nucleus (Beck and Hurt, 2016). The NPC also plays other roles in transcription and gene regulation, cytoskeletal regulation, and nuclear membrane architecture (Strambio-De-Castillia et al., 2010). During mammalian cell division, the NPC disassembles during open mitosis/meiosis and reassembles upon completion of chromosome segregation (Otsuka and Ellenberg, 2018). Emerging work demonstrates that the Nups, despite NPC disassembly, are not passive during cell division. Many Nups have roles during cell division including the promotion of microtubule nucleation, regulation of the anaphase promoting complex/cyclosome, and localization of spindle assembly checkpoint proteins Mad1 and Mad2 (Mossaid and Fahrenkrog, 2015; Garcia et al., 2021). For example, during interphase, Nup153 is a nuclear pore complex protein in the nuclear basket that plays roles in nuclear trafficking (Ball and Ullman, 2005) and transcription (Toda et al., 2017), but during mitosis, Nup153 plays roles in nuclear pore complex assembly and nuclear envelope modeling (Duheron et al., 2014; Vollmer et al., 2015) and spindle assembly checkpoint regulation via an interaction with Mad1 (Lussi et al., 2010; Mossaid et al., 2020).

Here, we identify Nup153 as a novel SPOP substrate. We demonstrate that SPOP binds, ubiquitylates, and degrades Nup153. We also demonstrate that SPOP overexpression results in the mislocalization of the spindle assembly checkpoint protein Mad1 from the nuclear envelope, presumably via depletion of Nup153. Altogether, our results demonstrate that Nup153 is a substrate of SPOP and suggest that the SPOP-Nup153 axis may be a mechanism by which SPOP regulates aspects of cell cycle progression and cell division.

## Methods

### Molecular cloning

To generate pDONR221 vectors containing the coding sequence (or truncated coding sequence) of each protein used in this study, primers containing Gateway cloning sites were designed against the coding sequence (or the desired truncation) of each gene and the genes were amplified via PCR. The PCR products were extracted and purified from an agarose gel. The appropriate pDONR221 plasmids and subsequent destination vectors were generated with Gateway cloning as previously described (Torres et al., 2009).

To generate amino acid substitution and deletion mutants, primers were designed with Agilent’s QuikChange Primer Design program and the mutagenesis was performed with Agilent QuikChange Lightning using the appropriate pDONR as a template according to manufacturer’s instructions. See Supplementary File 1 for all plasmids used in this study.

### Cell culture, drug treatments, and transfections

Cells were maintained in DMEM/F12 media supplemented with 10% FBS (v/v) and penicillin/streptomycin at 37°C in 5% CO_2_. HeLa and HEK293T cells were maintained with standard FBS, and Flp-In T-REx cells were maintained with FBS that contained no detectable tetracycline. The doxycycline-inducible Flp-In T-REx cell lines were generated as previously described (Bradley et al., 2016). The doxycycline-inducible protein was induced in Flp-In T-REx cells by addition of 0.1 μg/mL doxycycline for 16-24 hours. Cells were discarded within 12 weeks of thawing to avoid high passage numbers.

For plasmid DNA and siRNA transfections, for one well of a 12-well plate (surface area 3.5 cm^2^), about 80,000 cells were plated. The next day, the cells (at about 70% confluence) were transfected. For plasmid DNA transfection, 1 μg of total DNA was transfected with 4 μL of Fugene HD in 100 μL of Opti-MEM, according to manufacturer’s instructions. For control plasmid DNA transfections, non-coding DNA (pCS2-Myc backbone plasmid) was used. For siRNA transfection, 0.3 μL of 100 μM siRNA was transfected with 5 μL of Lipofectamine RNAiMax in 100 μL of Opti-MEM, according to manufacturer’s instructions (final concentration of siRNA: 30 nM). For control siRNA transfections, siGLO was used. The media was changed about 6-8 hours after plasmid DNA transfection or the next day (about 20 hours) after siRNA transfection. DNA transfected cells were assayed after 48 hours, while siRNA transfected cells were assayed after 72 hours.

For cell cycle arrests, cells were treated with 2 mM thymidine or 232 nM Taxol for 16-20 hours. For protein degradation experiments, cells were transfected with indicated plasmids, then, 48 hours after transfection, treated with 20 μM MG132, 50 mM chloroquine, or both for 5 hours. For in cell ubiquitylation experiments, pgLAP1-Nup153 HeLa cells were treated with 20 μM MG132 for 4-5 hours before lysis.

### Mass spectrometry

pgLAP2-SPOP^WT^ HEK293 Flp-In T-Rex cells were generated and tandem affinity purification was performed as previously described (Torres et al., 2009). The eluates were run onto an SDS-PAGE gel and ten gel slices were subjected for in-gel digestion with trypsin. Mass spectrometry and sequence analysis was performed at the Harvard Microchemistry and Proteomics Analysis Facility by microcapillary reverse-phase HPLC nano-electrospray tandem mass spectrometry (µLC/MS/MS) on a Thermo LTQ-Orbitrap mass spectrometer as previously described (Taniguchi et al., 2002).

### Cell lysis, co-immunoprecipitations, SDS-PAGE, and immunoblotting

To produce cell lysates, the media was removed and cells were lifted from the plate using cell dissociation solution (5% (v/v) glycerol, 1 mM EDTA, 1 mM EGTA in PBS) for 5 minutes at 37°C. The cells were pelleted by centrifugation at 500 x g for 5 minutes. Cell lysates, co-immunoprecipitations, SDS-PAGE, and immunoblotting were performed as previously described with the following modifications (Cheung et al., 2016):

1. The salt and detergent concentration in the lysis buffer was increased to 200 mM KCl and 1% (v/v) NP40. ATP was not added to the lysis buffer.
2. Cells were lysed on ice by vortexing for 3 seconds every 3 minutes, 5 times total, before clearing via centrifugation at 13.1k x g at 4°C for 10 minutes.
3. For co-immunoprecipitations, the lysis buffer was supplemented with 10 μM of MG132, 10 mM of NEM, and phosphatase inhibitor cocktail. Two volumes of lysis buffer without NP40 was added to the cell lysate during the binding so that the final concentration of NP40 during binding was 0.33%. Lysates were incubated with solid phase (either S-protein agarose, magnetic HA beads, or magnetic FLAG beads) for 2 hours at 4°C on a rotating platform. The solid phase was washed with the supplemented lysis buffer containing 0.2% NP40.
4. For co-immunoprecipitation experiments using S-Tag agarose, 200 μL S-Tag agarose slurry was used (about 100 μL packed volume). The agarose solid phase was collected by centrifugation at 500 x g for 2 minutes at 4°C.
5. The lysates were run on 4-20% 15-well SDS-PAGE gels.

For immunoblots, molecular weight markers are shown to the right in kDa. Blots were imaged on a LI-COR Odyssey fluorescence scanner. In some cases, for a given experiment, lysates were loaded equivalently onto multiple gels and/or the blot was cut horizontally to allow for probing by multiple antibodies. See Supplementary File 1 for antibodies used and their dilutions.

### *In vitro* transcription/translation and binding experiments

To produce IVT-expressed protein, 200 ng of indicated pCS2 plasmid was added to 8 μL of SP6 TNT-master mix, 0.5 μL of 1 mM methionine, and the volume was brought to 10 μL per reaction. The tubes were incubated at 30°C for 1.5 hours with shaking at 250 rpm. Upon completion of the incubation, 0.5 μL (approximately 5%) of the protein was saved for the input portion of a gel and heated at 95°C for 5 minutes with 3 μL of water and 5 μL of 2x Laemmli buffer. The binding reaction, washing, and elution was performed in the same manner as detailed for co-immunoprecipitations, except the concentration of NP40 was 0.25% throughout.

### Immunofluorescence and microscopy

For fixed cell immunofluorescence microscopy, HeLa cells were plated onto poly-D-lysine coated coverslips in a 24-well plate. The same transfection procedures as a 12-well plate were followed, except everything was halved to adjust for the smaller surface area. Fixation with PFA, staining, and microscopy were performed as previously described (Guo et al., 2021). Methanol fixation was performed via the addition of absolute methanol cooled to −20°C and fixing the cells at −20°C for 20 minutes. See Supplementary File 1 for antibodies used and their dilutions.

Images were acquired as a single plane or as a z-stack of 8 images encompassing a z-volume of 6.23 microns. For z-stack images, the maximum projection of the images is presented. Figures were processed in FIJI and exported as .tif files.

### Live cell imaging

HeLa cells were plated in glass-bottom 24-well plates and transfected with plasmid DNA. Twenty-four hours after transfection, the cells were arrested with 2 mM thymidine. Twenty-four hours after thymidine addition, the cells were released from thymidine arrest via two washes with PBS and two washes with media. Five hours after thymidine release (that is, 53 hours after DNA transfection), the cells were transferred to the imaging system. Within the imaging system, the cells were maintained in 5% CO_2_ in 20% oxygen and 75% nitrogen and at 37°C.

Images were acquired on a LSM880 platform at 1024×1024 pixel resolution with a Plan-Apochromat 10x/0.45 NA M27 air objective. Images were acquired at 10-minute intervals for 10 hours. For the GFP channel, an argon laser (excitation 488 nm, emission window 493-578 nm) and, for transmitted light (bright field), a DPSS 561-10 laser (excitation 561 nm, emission window 586-696 nm) were used. A z-stack of 6 images encompassing a z-volume of 6.12 microns was acquired, and this z-stack was used to generate the maximum projection image.

### CellProfiler quantification and other code

A Python script used to identify SPOP consensus binding sites is available on Github: https://github.com/jong2ucla/PythonScripts/blob/main/SPOP_ConsensusFinder.

The CellProfiler pipeline used to quantify nuclear envelope-localized Nup153 or MAD1 fluorescence to DNA fluorescence is available on Github: https://github.com/jong2ucla/CellProfiler/blob/main/rbNup153toDNA.cppipe.

### RT-qPCR

For RNA extraction, between 70-72 hours after siRNA transfection, the media was removed from HeLa cells and 200 μL of TriReagent was added directly to each well of the plate to lyse the cells. RNA extraction, Dnase I digestion, RT-PCR, and qPCR were performed as previously described (Guo et al., 2021). For RT reactions, 750-1500 ng of RNA was used, and for qPCR reactions, 75-150 ng of cDNA with 400 nM each of forward and reverse qPCR primer was used.

Data were processed using the delta delta Ct method as previously described, using Gapdh as a housekeeping gene for normalization (Livak and Schmittgen, 2001). qPCR values were normalized to cells transfected with control siGLO.

### Statistical analyses

For image quantification, statistical significance was determined via one-way ANOVA and Tukey-HSD. For qPCR quantification, each siRNA transfection, RNA extraction, and RT-PCR were performed three separate times (three biological replicates). For each biological replicate, the cDNA was assayed twice via qPCR (two technical replicates). For reported knockdown efficiencies, the two technical replicates of a biological replicate were averaged, and these averaged values were used to calculate the reported average and standard deviation (avg±SD) of the three biological replicates. Statistical significance was determined via one-way ANOVA and Tukey-HSD.

## STAR Protocols

| REAGENT or RESOURCE | SOURCE | IDENTIFIER |
| --- | --- | --- |
| **Recombinant DNA** |  |  |
| See Supplementary File 1 for all plasmids and vectors used in this study. |  |  |
| **Oligonucleotides** |  |  |
| SPOP siRNA #100 | Millipore Sigma | siRNA ID:<br>SASI_Hs01_00034100 |
| SPOP siRNA #101 | Millipore Sigma | siRNA ID:<br>SASI_Hs01_00034101 |
| SPOP siRNA #102 | Millipore Sigma | siRNA ID:<br>SASI_Hs01_00034102 |
| siGLO Red Transfection Indicator | Dharmacon | D-001630-02-05 |
| Oligonucleotide sequences for siRNA, qPCR primers, and primers used for cloning are in Supplementary File 1 | Eurofins Genomics | See Supplementary File 1 |
| **Experimental Models: Cell Lines** |  |  |
| HeLa Flp-In T-REx | Stephen Taylor (University of Manchester) | PMID: 18541701 (Tighe et al., 2008) |
| pgLAP1-Nup153 HeLa Flp-In T-REx | This paper |  |
| Flp-In™ T-REx™ 293 Cell Line | Invitrogen | R78007 |
| pgLAP2-SPOP<sup>WT</sup> HEK293 Flp-In T-REx | This paper |  |
| HeLa | ATCC | CCL-2 |
| HEK293T | ATCC | CRL-3216 |
| **Antibodies** |  |  |
| See Supplementary File 1 for all primary and secondary antibodies used for immunoblotting and immunofluorescence microscopy in this study. |  |  |
| **Critical Commercial Assays** |  |  |
| QuikChange Lightning Site-Directed Mutagenesis Kit | Agilent | 210518 |
| Fetal Bovine Serum – Premium | Atlanta Biologicals | Cat No: S10350, Lot No: F17085 |
| 4–20% Mini-PROTEAN® TGX™ Precast Protein Gels, 15-well, 15 µL | BIORAD Life Science Group | #4561096 |
| Precision Plus Protein Dual Color Standards, 500 µL | BIORAD Life Science Group | #1610374 |
| Fetal Bovine Serum - Premium, USDA Origin Tetracycline Free | BIOWEST LLC | Cat No: S162TA, Lot No: 045G19 |
| Gibco™ Opti-MEM™ I Reduced Serum Medium | Fisher Scientific | 31-985-070 |
| MilliporeSigma™ Immobilon-FL PVDF Membrane | Fisher Scientific Co. | IPFL00005 |
| qPCRBIO SyGreen Blue Mix Separate-ROX | Genesee Scientific | 17-507 |
| UltraScript 2.0 cDNA Synthesis Kit | Genesee Scientific | 17-702 |
| DMEM/Ham's F-12, with L-Glutamine | Genesee Scientific | 25-503 |
| Trypsin-EDTA, 0.25% 1X, phenol red | Genesee Scientific | 25-510 |
| BP Clonase II | Invitrogen | 11789020 |
| LR Clonase II | Invitrogen | 11791020 |
| Anti-HA-tag mAb-Magnetic Beads (Monoclonal Antibody) | MBL International | M132-11 |
| Phosphatase Inhibitor Cocktail 2 | Millipore Sigma | P5726-1ML |
| TNT® SP6 Quick Coupled Transcription/Translation System | Promega | L2080 |
| FuGENE® HD Transfection Reagent, 1ML | Promega | E2311 |
| QIAquick DNA gel extraction kit | Qiagen | 28706 |
| Anti-FLAG® M2 Magnetic Beads | Sigma-Aldrich Corp. | M8823-1ML |
| Platinum Taq DNA Polymerase | Thermo Fisher Scientific | 10966018 |
| Pierce™ Protease Inhibitor Tablets, EDTA-free | Thermo Fisher Scientific | A32965 |
| Tris-Glycine-SDS, 10X Solution (Electrophoresis), Fisher BioReagents | Thermo Fisher Scientific | BP13414 |
| PBS, Phosphate Buffered Saline, 10X Solution, Fisher BioReagents | Thermo Fisher Scientific | BP39920 |
| TAE Buffer, Tris-Acetate-EDTA, 50X Solution, Electrophoresis, Fisher BioReagents | Thermo Fisher Scientific | BP13324 |
| Lipofectamine™ RNAiMAX Transfection Reagent | Thermo Fisher Scientific | 13778150 |
| Invitrogen™ Molecular Probes™ ProLong™ Gold Antifade Mountant | Thermo Fisher Scientific | P36934 |
| EMD Millipore™ Novagen™ S-protein Agarose | Thermo Fisher Scientific | Novagen 69704-4 |
| **Hardware and Software** |  |  |
| S1000 Thermal Cycler with Dual 48/48 Fast Reaction Module | BIORAD Life Science Group | 1852148 |
| CFX Connect Real-Time PCR Detection System; software: CFX Manager, Version 3.0 | BIORAD Life Science Group | 1855201 |
| CellProfiler v4.2.1 | https://cellprofiler.org/ | PMID: 34507520 (Stirling et al., 2021) |
| ImageJ2/FIJI | https://imagej.net/software/fiji/ | PMID: 22743772 (Schindelin et al., 2012) |
| Leica DMI6000 microscope (Leica DFC360 FX Camera, 63x/1.40–0.60 NA oil objective); software: Leica AF6000 | Leica |  |
| LI-COR Odyssey Imager; software: Odyssey v3.0 | LI-COR | Model 9120 |
| NanoDrop 2000c; software: NanoDrop 2000/2000c version 1.6.198 | Thermo Fisher Scientific | ND-2000C |
| SuperPlots of Data | https://huygens.science.uva.nl/SuperPlotsOfData/ | PMID: 33476183 (Goedhart, 2021) |
| Inkscape 0.92.4 | https://inkscape.org/ |  |
| LSM880 microscope; software: ZEN Black (v2.3) | Carl Zeiss Microscopy | PMID: 34813726 (Markaki et al., 2021) |

## Results

### SPOP binds Nup153

To identify novel substrates of SPOP, we overexpressed tagged SPOP in HEK293 cells and analyzed tandem affinity-purified complexes via mass spectrometry (Supplementary Figure 1A). Our results showed a number of known SPOP substrates and interactors, such as Caprin1 (Shi et al., 2019) and G3BP1 (Mukhopadhyay et al., 2021). The most abundant protein identified was Nup153, a nuclear pore complex protein, and its binding partners Nup50 and some importin subunits like KPNA6. A direct interaction between these proteins is involved in nuclear trafficking through the nuclear pore complex (Makise et al., 2012). Given that Nup153 (Lan et al., 2019), Nup50 (Hjorth-Jensen et al., 2018), KPNA6 (Ewing et al., 2007; Yuan et al., 2020), and other importin subunits (Huttlin et al., 2021; Yuan et al., 2020) have all been identified as possible interactors of SPOP by mass spectrometry, and given their high abundance in our mass spectrometry results, we pursued these proteins as possible substrates of SPOP.

To determine whether SPOP binds these substrates, we expressed HA-tagged SPOP^WT^, its binding mutant SPOP^F102C^, and FLAG-tagged Nup153, Nup50, and KPNA6 in a cell-free, *in vitro* transcription and translation rabbit reticulocyte lysate system (hereafter, IVT-expressed protein). We also expressed FLAG-tagged Cdc20, a known SPOP substrate (Wu et al., 2017), as a positive control, and Plk1, which lacks SPOP binding consensus motifs (Zhuang et al., 2009), as a negative control. Binding reactions between these SPOP constructs and potential substrates demonstrated that SPOP^WT^ could readily immunoprecipitate the positive control Cdc20, Nup153, Nup50, and KPNA6, and the binding mutant SPOP^F102C^ could only weakly immunoprecipitate KPNA6 (Figure 1A).

**Figure 1.**
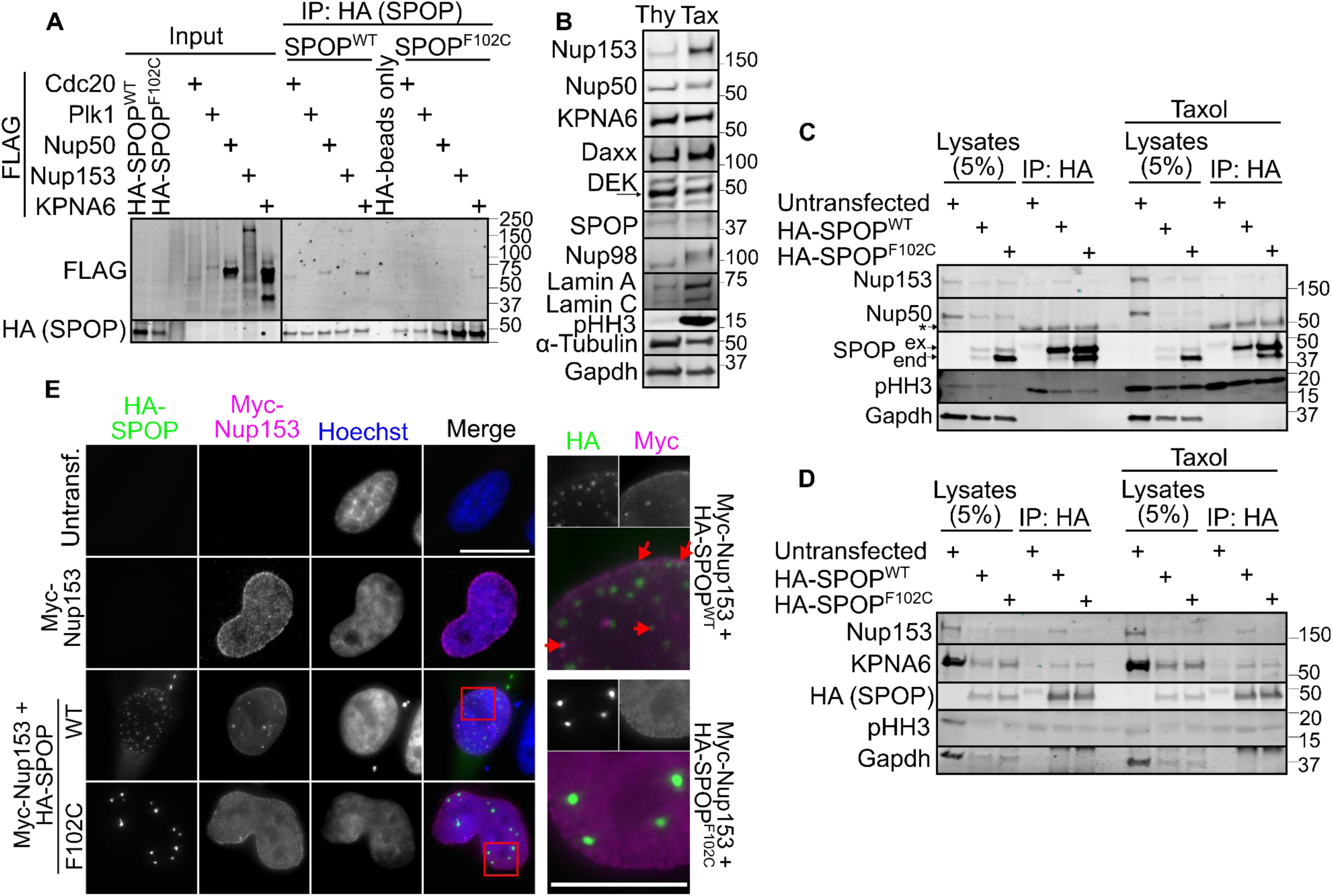
SPOP binds to and colocalizes with Nup153. (A) IVT-expressed HA-SPOP and FLAG-substrates underwent HA-immunoprecipitation to determine which substrates can bind to SPOP. Cdc20 was used as a known SPOP substrate (positive control) and Plk1 as a negative control. (B) HeLa cells were arrested with 2 mM thymidine (Thy) or 232 nM Taxol (Tax) for 17 hours and the lysates were probed for the indicated proteins. pHH3 serves as a mitotic marker. (C, D) HeLa cells were transfected with HA-SPOP and either left asynchronous or treated with 232 nM Taxol. The lysates were subjected to HA-immunoprecipitation and the resulting blots were probed with the indicated antibodies. In (C), *ex* refers to exogenous (HA-tagged) and *end* refers to endogenous SPOP. (E) HeLa cells transfected with Myc-Nup153 and HA-SPOP were imaged after staining with the indicated antibodies. The scale bars in both the full images and the insets are 10 μm.

Having established that SPOP and these potential substrates can associate as proteins expressed in a cell-free system, we sought to establish if these proteins associate in human cell lysates via co-immunoprecipitation experiments. We first observed that Nup153 and known SPOP substrates Caprin1 and Myd88 were able to co-immunoprecipitate with exogenous HA-SPOP^WT^ but not the binding mutant HA-SPOP^F102C^ (Supplementary Figure 1B). We also observed that Nup153 levels, but not Nup50 levels, increased in cells arrested in mitosis via taxol treatment relative to cells arrested in G1/S phase via thymidine treatment (Figure 1B). Such a result is consistent with previous reports that some nucleoporins, including Nup153, generally increase from G1 to G2/M (Chakraborty et al., 2008), coinciding with nuclear pore complex synthesis and assembly during S phase. Another SPOP substrate, DAXX, also increased in taxol-arrested cells relative to asynchronous cells (Kwon et al., 2006); in contrast, the levels of another SPOP substrate, DEK, did not have a cell cycle phase dependence (Theurillat et al., 2014), suggesting that cell cycle-dependent protein levels are not shared among all SPOP substrates.

We wondered then if SPOP regulation of the substrates we identified could be differentially regulated in mitosis. This question was supported by two opposing reasons: first, SPOP-substrate interactions have been known to be enhanced by or dependent on substrate phosphorylation (Wang et al., 2020a, 2021; Gan et al., 2015; Li et al., 2011; Jiang et al., 2021) (although phosphorylation of some substrates weaken SPOP-substrate binding (Ostertag et al., 2019b; Zhuang et al., 2009; Zhang et al., 2009; Wang et al., 2019)). Nuclear pore complex proteins are phosphorylated by mitotic kinases as the cell enters into mitosis, an event that precedes nuclear envelope breakdown and the dissociation of the nuclear pore complex (Linder et al., 2017; Martino et al., 2017). Indeed, we observed a significant decrease in the mobility of Nup98 in the mitotic population, suggestive of post-translational modifications such as phosphorylation (Figure 1B). While not as dramatic, we also observed a slight decrease in mobility for Nup153 and Nup50 in the mitotic population as well, suggestive of post-translational modifications such as phosphorylation, a modification which has been previously documented (Ball and Ullman, 2005). Whether or not these mitotic phosphorylation events would promote or decrease SPOP binding to our substrates was an open question.

On the other hand, during mitosis, the nuclear envelope is disassembled and SPOP can no longer associate with the nuclear pore complex proteins. Dilution of SPOP into the cytoplasm could weaken the SPOP-nuclear pore complex protein interaction. In a similar vein, SPOP was previously reported to be active in interphase and deactivated during late mitosis due to degradation by the APC/C substrate adaptor Cdh1/Fzr1 (Zhang et al., 2017a). We note that whereas Zhang et al. observed a decrease of SPOP protein levels in mitosis and an increase of SPOP protein levels in interphase via immunoblotting, we observe no difference in SPOP levels between the two phases (Figure 1B). Whether or not inactivation of SPOP during mitosis, either due to degradation or dilution (or some other mechanism), was also an open question.

To assess whether mitotic events could mediate the effect of SPOP binding to our identified substrates, we performed co-immunoprecipitation experiments with asynchronous, cycling cells (about 80% in G1/S) and taxol-arrested mitotic cells (about 80% in M phase) (Figure 1 C, D). We observed that, similar to our previous results, Nup153 co-immunoprecipitates with SPOP in the asynchronous cell population. However, the amount of immunoprecipitated Nup153 was less in the mitotic cell population than in the asynchronous population, suggesting that the SPOP-Nup153 interaction is weaker in mitotic cells (when Nup153 is hyperphosphorylated and the nuclear envelope has broken down) than in interphase cells (which contain an intact nuclear envelope).

Despite observing a binding interaction between SPOP and Nup50 with IVT-expressed proteins, we did not observe co-immunoprecipitation of Nup50 with SPOP in either an asynchronous or mitotic cell lysates (Figure 1C). This result may suggest that the SPOP-Nup50 interaction may be weak or transient within cells. Such an idea is supported by the highly mobile nature of Nup50 and its transient association with the nuclear pore complex (and thus perhaps SPOP) (Buchwalter et al., 2014). We also observed that KPNA6 co-immunoprecipitated with both SPOP^WT^ and its binding mutant SPOP^F102C^ in both the asynchronous and mitotic cell lysates (Figure 1D). Why the mutant construct of SPOP can bind KPNA6 in cell lysates but not with IVT-expressed protein is unclear. One possibility is that in cell lysates, endogenous SPOP can oligomerize with the mutant SPOP such that the interaction between exogenous SPOP^F102C^ and endogenous KPNA6 is mediated through endogenous, wild-type SPOP (Pierce et al., 2016) (also note the stronger co-immunoprecipitation of endogenous SPOP with exogenous HA-SPOP^F102C^ than HA-SPOP^WT^ in Figure 1C). Consistent with this idea, we also observe that, in HEK293T cells, expression of SPOP^F102C^ also results in weaker (relative to SPOP^WT^) but detectable co-immunoprecipitation of Nup153 and KPNA6 (Supplementary Figure 1C).

Because Nup153 presented the clearest case of SPOP binding (that is, we observed the SPOP-Nup153 interaction in both IVT-expressed proteins and in HeLa cell lysates, and the interaction was disrupted by the canonical SPOP binding mutant F102C), we proceeded to examine the possibility of Nup153 as an SPOP substrate. We first sought to determine if the two proteins colocalize in the nucleus by transiently transfecting DNA coding for both proteins into HeLa cells. We observed that SPOP^WT^ and Nup153 colocalized along the nuclear envelope and in some nuclear speckles, similar to the localization observed with SPOP and other SPOP substrates (Bunce et al., 2008; Marzahn et al., 2016). In contrast, the binding mutant SPOP^F102C^ had a different localization, and these nuclear speckles exclude Nup153. In particular, Nup153 lacked nuclear speckle localization when co-expressed with SPOP^F102C^, suggesting that Nup153 associates with SPOP^WT^ but not SPOP^F102C^ compartments, similar to what has been observed previously for other SPOP-substrates coacervates (Bouchard et al., 2018).

Altogether, these results demonstrate SPOP binds to and colocalizes with Nup153, suggesting that Nup153 is a substrate of SPOP.

### SPOP binds to Nup153 at least within the N-terminal Nuclear Pore Complex domain

The SPOP MATH domain binds to an SPOP binding consensus site on its substrates, a canonical five amino acid motif of nonpolar–polar–S–S/T–S/T, though some motifs with mismatches (underlined) are known, such as with GL9 (Zhang et al., 2021) (GL9 SPOP site #1: AD<u>T</u>TS; GL9 SPOP site #2: AD<u>T</u>TT) and Myd88 (Guillamot et al., 2019; Jin et al., 2019a) (VDSS<u>V</u>). One SPOP substrate, Pdx1, has SPOP binding consensus sites that are significantly different from the canonical SPOP binding consensus site (Pdx1 SPOP site #1 (Ostertag et al., 2019b): VTS<u>GE</u>; Pdx1 SPOP site #2 (Usher et al., 2021): PQ<u>P</u>SS) but nonetheless use the same binding modality in the MATH domain. Therefore, we sought to determine the SPOP binding consensus sites on Nup153.

Unfortunately, unlike other SPOP substrates which tend to have only a few SPOP binding consensus sites, Nup153 contains 15 SPOP canonical binding consensus motifs (and many more “near-miss” binding sites where a mismatch is allowed). Individually mutating these sites was deemed impractical. Instead, we opted to first truncate Nup153 to determine which domains could bind to SPOP, and then mutate the SPOP binding consensus motifs of the Nup153 domains that bound to SPOP. To do this, Nup153 was truncated according to its three domains: the N-terminal nuclear pore complex domain (NPC), the central zinc finger domain (ZnF), and the C-terminal phenylalanine and glycine repeat domains (FG). We also generated the NPCZnF and ZnFFG domains (Figure 2A). Immunoprecipitation experiments using IVT-expressed proteins demonstrated that SPOP binds to the Nup153 NPC, NPCZnF, ZnFFG, and FG truncations (Figure 2B). Altogether, these results suggest that SPOP binds to the Nup153 NPC and the FG domains and not the ZnF domain.

**Figure 2.**
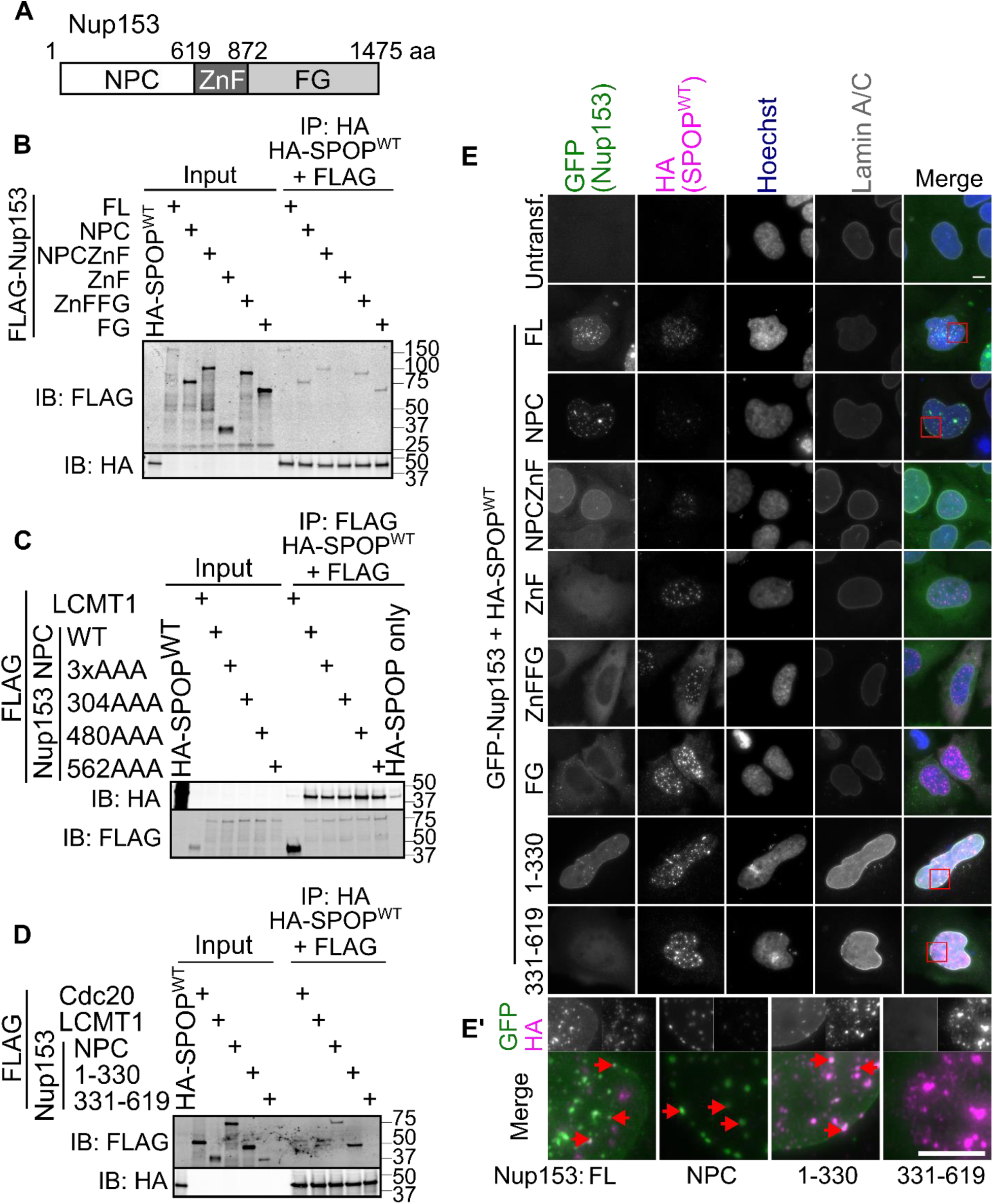
SPOP binds at least to the N-terminal NPC domain of Nup153. (A) Nup153 was truncated according to its domains. NPC, nuclear pore complex domain; ZnF, zinc finger; FG, Phe-Gly repeat domain. (B) IVT-expressed HA-SPOP and FLAG-Nup153 truncations were subjected to HA-immunoprecipitation and the resulting blots probed with HA and FLAG antibodies. (C) IVT-expressed HA-SPOP and FLAG-Nup153 NPC with indicated alanine substitutions were subjected to FLAG-immunoprecipitation and the resulting blots probed with HA and FLAG antibodies. FLAG-LCMT1 served as a negative control. SPOP only refers to a condition with HA-SPOP and the FLAG beads (no FLAG-tagged protein). (D) Same as (C), except with FLAG-Nup153 NPC, 1-330, or 331-619. FLAG-Cdc20 is used as a known SPOP substrate (positive control). (E) HeLa cells transfected with (GFP) pgLAP1-Nup153 truncations and HA-SPOP were imaged (z-stack) after staining with the indicated antibodies. (E’) Insets showing colocalization of HA-SPOP and GFP-Nup153. The scale bars in both the full images and the insets are 10 μm.

To further verify that SPOP could bind to the NPC and FG domains of Nup153, we co-expressed each Nup153 truncation with SPOP^WT^ in HeLa cells. We observed co-localization of SPOP and Nup153 NPC and NPCZnF, but not any of the other truncations (Figure 2E and E’). We note that the truncations of Nup153 that lack the N-terminal NPC domain did not localize within the nucleus and thus would not bind to the nuclear protein SPOP (Ball and Ullman, 2005). The mislocalization of these Nup153 truncations that do not contain the N-terminal NPC domain may bias the interpretation of this experiment, at least for these truncations. In particular, the observation that SPOP does not co-localize with the Nup153 FG domain does not rule out the possibility that SPOP may bind to the FG domain of Nup153. Nonetheless, these results suggest that the Nup153 NPC domain co-localized with SPOP, further suggesting that SPOP binds to Nup153 at least within the N-terminal NPC domain. Given the easy read-out that we could obtain via microscopy, we opted to continue characterization of the Nup153 NPC domain instead of the FG domain.

The Nup153 NPC domain contains three SPOP binding consensus sites at amino acids ^303^V<u>TSS</u>T^308^, ^479^I<u>TSS</u>S^484^, and ^561^G<u>SSS</u>T^566^. To determine which binding site(s) were sites of SPOP-Nup153 NPC binding, we mutated the middle three amino acids (underlined in the previous sentence) to three alanine residues, generating Nup153 NPC 304AAA, 480AAA, and 562AAA, an approach adopted from studies of other SPOP substrates like Geminin (Ma et al., 2021) and GLP (Zhang et al., 2021). We also generated a construct of the Nup153 NPC whereby all three aforementioned sites were mutated to AAA (3xAAA). LCMT1, a protein that contains no SPOP consensus binding sites, was used as a negative control (Xia et al., 2015).

Surprisingly, all four alanine constructs of the Nup153 NPC, including the 3xAAA where all three SPOP binding consensus sites were mutated, still bound to SPOP (Figure 2C). Intrigued, we again truncated the Nup153 NPC into two truncations, chosen to avoid disrupting secondary structures within the NPC: NPC amino acids 1-330 and NPC amino acids 331-619. Binding experiments with IVT-expressed proteins demonstrated that the Nup153 NPC 1-330 but not 331-619 binds to SPOP (Figure 2D). Via microscopy, the Nup153 NPC 1-330 domain localized within the nucleus but Nup153 NPC 331-619 did not. Similar to how the Nup153 FG domain does not localize within the nucleus, we were also unable to use the localization of the Nup153 NPC 331-619 to support an SPOP binding site. Nonetheless, given the binding of Nup153 NPC 1-330 to and colocalization with SPOP, we pursued the SPOP binding site within the first 330 amino acids of Nup153.

### SPOP binding to the Nup153 NPC is not dependent on traditional SPOP binding consensus motifs

To determine where the SPOP-Nup153 1-330 binding site is located, we again mutated Nup153 1-330 by introducing premature stop codons about every 30 amino acids (with some consideration to not disrupt predicted secondary structure) to generate C-terminally truncated constructs of Nup153 that spanned from the full 1-330 amino acids to the smallest segment generated, 1-167 amino acids. Binding reactions with these IVT-expressed proteins demonstrated that SPOP binds only to the full 1-330 amino acids, as Nup153 1-300 and smaller constructs exhibited weaker binding to SPOP (Supplementary Figure 2A). We thus further examined these final 30 amino acids in detail. In these 30 amino acids (^301^SYG<u>VTSST</u>ARRI<u>LQSLE</u>KM<u>SSPLAD</u>AKRIPS^330^), we identified three polar amino acid stretches (underlined in this sentence) that could serve as potential SPOP-Nup153 1-330 binding sites: amino acids 303-307 (the exact match to SPOP binding consensus motifs previously mutated to AAA), amino acids 312-316 (which resembles the first Pdx1 SPOP consensus binding site VTSGE as both sites contain a mismatch in the fourth position and a glutamate residue in the fifth position), and amino acids 319-214 (which we selected for its prevalence of serine residues).

We deleted these amino acids to generate Nup153 NPC 1-330 Δ303-307, Nup153 NPC 1-330 Δ312-316, and Nup153 NPC 1-330 Δ319-324. Surprisingly, mutation of any one of these sites did not disrupt the Nup153 NPC 1-330 construct from binding to SPOP (Supplementary Figure 2B). We confirmed that binding of Nup153 NPC 1-330 was dependent on the substrate-binding MATH domain of SPOP by performing the same experiment but with the SPOP mutant F102C (Supplementary Figure 2C). Combining two of these deletions in the Nup153 1-330 also did not disrupt Nup153 1-330 binding to SPOP, as the double deletion constructs (denoted as Δ1+Δ2, Δ1+Δ3, and Δ2+Δ3 for which deletions were generated) also bound to SPOP with the same affinity as the intact Nup153 1-330 construct (Supplementary Figure 2D). These results suggested that unlike other SPOP substrate binding interfaces, which can be easily disrupted by deleting or mutating approximately 5 amino acids, the SPOP-Nup153 binding interface within the Nup153 1-330 may span over a larger stretch of amino acids and/or may involve structural motifs in three-dimensional space that are not disrupted by deletions of small portions of the substrate.

Consistent with this idea, deleting the same three regions either individually or simultaneously within the entire Nup153 NPC (amino acids 1-619) did not disrupt SPOP-Nup153 binding (Supplementary Figure 2E). This result suggests that other amino acids within Nup153 residues 331-619 can compensate for the lack of the potential SPOP binding site within Nup153 residues 300-330. One hypothesis is that SPOP forms oligomers that can bind to multiple binding motifs on its substrate (Pierce et al., 2016). Consequently, an oligomeric chain of SPOP may bind to the Nup153 NPC (or full length Nup153) at multiple locations, granting an interaction with higher avidity. An additional SPOP binding site(s) – particularly a non-canonical binding site that does not match the usual consensus motif – within the Nup153 NPC may explain why deletion of residues 300-330 within the Nup153 NPC does not abrogate binding to SPOP. Altogether, these results suggest that the SPOP-Nup153 NPC binding interface is not dependent on canonical SPOP consensus binding sites and is complex.

### SPOP targets Nup153 for ubiquitin-mediated degradation

Having established that SPOP bound to Nup153, we sought to establish the consequence of this interaction: namely, does SPOP target Nup153 for ubiquitin-mediated degradation? Most SPOP substrates are degraded (Wang et al., 2020b; Cuneo and Mittag, 2019), but some, such as MacroH2A (Hernández-Muñoz et al., 2005), HIPK2 (Jin et al., 2021), and G3PB1 (Mukhopadhyay et al., 2021), are not. We began by overexpressing SPOP WT or its binding mutant F102C and probing the substrates of SPOP by immunoblotting. We saw a decrease in Nup153 and KPNA6, but not Nup50, upon expression of SPOP^WT^ but not SPOP^F102C^ (Figure 3A). We saw a corresponding decrease in known SPOP substrates Caprin1 and Myd88 (immunoblots are quantified in Supplementary Figure 3A). We were also able to recapitulate these results for GFP-tagged constructs of SPOP: GFP-SPOP^WT^ and endometrial cancer-associated mutant GFP-SPOP^E50K^ (a mutation in the MATH domain which can enhance (Janouskova et al., 2017) or weaken (Gao et al., 2023) SPOP-substrate binding and degradation, depending on the substrate and the cell type), but not GFP only or GFP-SPOP^F102C^, similarly degraded Nup153 and KPNA6 (Supplementary Figure 3C, D). These results suggest that SPOP targets Nup153 and KPNA6, but not Nup50, for degradation. We note that the observed pattern for protein degradation is the same as our co-immunoprecipitation results (Figure 1C, D): that is, we neither detected co-immunoprecipitation of Nup50 by SPOP nor a change in Nup50 levels upon SPOP overexpression. Overexpression of HA-SPOP WT, but not the binding mutant F102C, also resulted in lower levels of Nup153 via immunofluorescence microscopy (Figure 3B, C).

**Figure 3.**
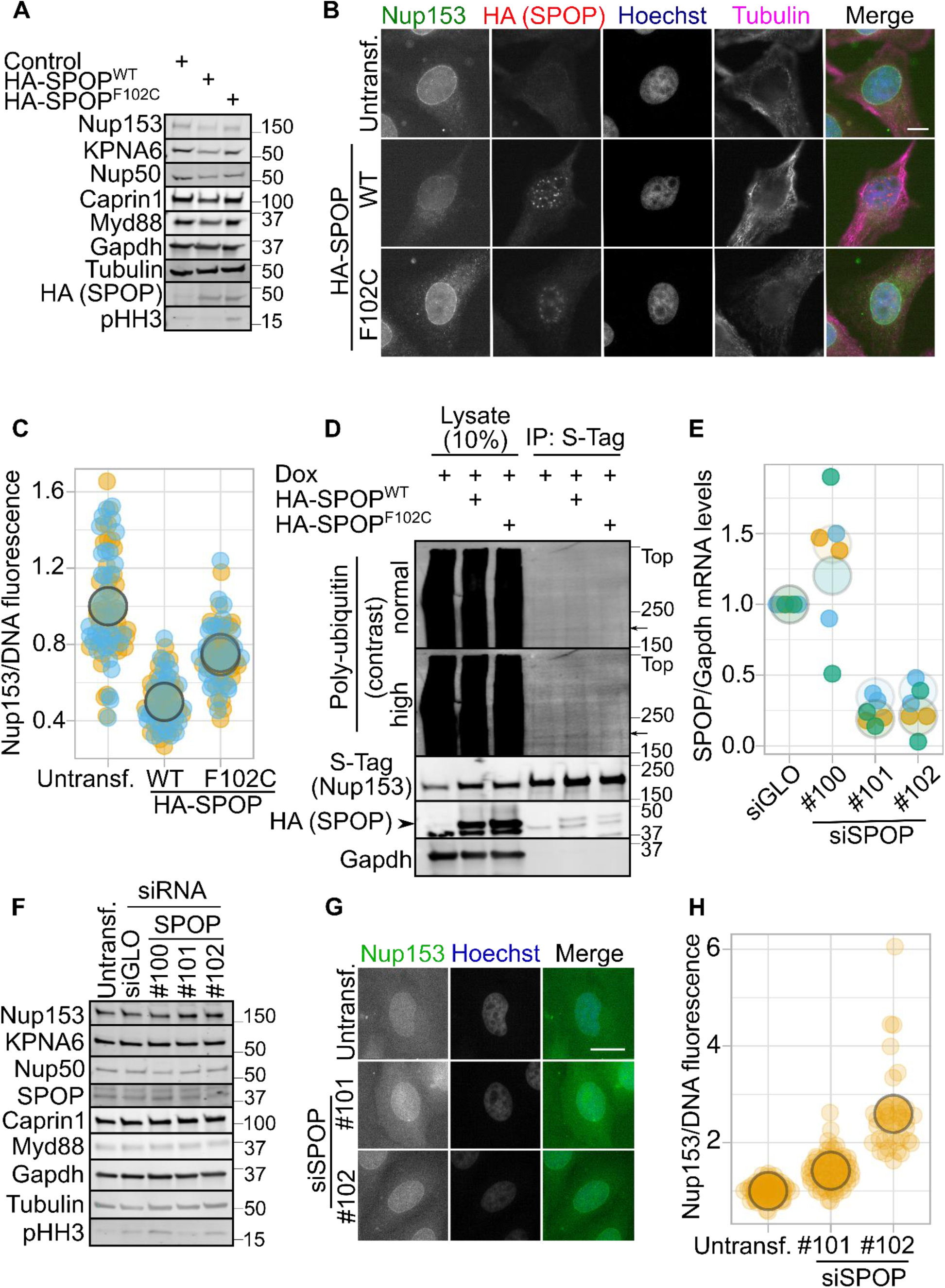
SPOP degrades Nup153. (A) Immunoblots from lysates of HeLa cells overexpressing HA-SPOP WT or F102C or non-coding DNA (control) were probed with the indicated antibodies. (B) HeLa cells transfected with HA-SPOP WT or F102C were fixed and stained with the indicated antibodies and (C) the ratio of the amount of nuclear envelope-localized Nup153 to DNA fluorescence was quantified. (D) pgLAP1-Nup153 HeLa cells were transfected HA-SPOP WT or F102C and pgLAP1-Nup153 expression was induced with doxycycline (dox). The lysates were subjected to S-Tag immunoprecipitation and probed with the indicated antibodies. The arrow indicates the location of pgLAP1-Nup153, and *Top* indicates the top of the gel. (E) mRNA levels of SPOP relative to Gapdh, normalized to siGLO, were determined by qPCR in HeLa cells transfected with the indicated siRNAs. Each color corresponds to a different biological replicate, and cDNA from each biological replicate was analyzed twice. Error bars represent the mean and standard deviation. p-values compared to siGLO: siSPOP #100: p=0.030; siSPOP #101: p<0.00001; siSPOP #102: p<0.00001. (F) Immunoblots from lysates from HeLa cells treated with the indicated siRNAs were probed with the indicated antibodies. (G) HeLa cells treated with the indicated siRNAs were fixed and stained with the indicated antibodies and the ratio of the amount of nuclear envelope-localized Nup153 to DNA fluorescence was quantified. The scale bar is 10 μm in (B) and 20 μm in (G). For (C), (E), and (H), a different color indicates a different biological replicate, individual measurements are shown as a small dot, and the average value of a biological replicate is presented as a large circle. For (A) and (F), the protein levels as determined by immunoblot is quantified in Supplementary Figure 3. Untransf., untransfected cells.

To determine the mechanism of Nup153 degradation, we again overexpressed SPOP WT or its binding mutant F102C and treated the cells with proteasomal inhibitor MG132, lysosomal inhibitor chloroquine (an inhibitor of autophagy (Mauthe et al., 2018)), both MG132 and chloroquine, or left the cells untreated. Both MG132 and chloroquine prevented SPOP^WT^-mediated degradation of Nup153, suggesting that Nup153 degradation may be both via proteasomal and lysosomal degradation (Supplementary Figure 3B).

Having determined that SPOP overexpression degrades Nup153, we sought to determine whether SPOP ubiquitylates Nup153. We generated a HeLa cell line that expressed GFP-S Tag-Nup153 in a doxycycline-inducible manner (Torres et al., 2009). We then transiently transfected either SPOP WT or its binding mutant F102C, induced GFP-S Tag-Nup153 overexpression via doxycycline addition to cell culture media, and precipitated the exogenously expressed Nup153. We probed immunoblots from these lysates with an antibody that detects polyubiquitin chains. These blots showed that polyubiquitylation of Nup153 is increased when SPOP^WT^, but not SPOP^F102C^, is overexpressed (Figure 3D), suggesting that the degradation of Nup153 by SPOP is mediated through ubiquitylation. However, whether this degradation is proteasomal or lysosomal (or both) is not clear.

We then determined the effect of SPOP knockdown on Nup153 levels. We tested three siRNAs against SPOP, siSPOP #100, #101, and #102. We assessed the ability of these siRNAs to knockdown SPOP by RT-qPCR (Figure 3E) and immunoblotting (Figure 3F and Supplementary Figure 3E).We were able to readily detect knockdown of SPOP mRNA levels via RT-qPCR (normalized to Gapdh mRNA levels and relative to control, non-targeting siRNA siGLO; Figure 3D) with siSPOP #101 (24±9% SPOP mRNA remaining) and #102 (27±10% SPOP mRNA remaining) but not siSPOP #100 (128±13% SPOP mRNA remaining). In HeLa cells, use of these siRNAs demonstrated that siSPOP #101 and #102, but not the SPOP siRNA #100 (the siRNA that did not knockdown SPOP) nor the control siGLO, resulted in an increase in Nup153 and Caprin1 protein levels and a modest increase in Myd88 levels (Figure 3E). No change in protein levels was detected for KPNA6 or Nup50. We note that protein levels of KPNA6, which co-immunoprecipitates with SPOP and is degraded upon SPOP overexpression, do not change upon SPOP knockdown. We are unsure as to whether the moderate change in Myd88 levels, a known SPOP substrate of degradation, and the apparent no change in protein levels for KPNA6 are due to an incomplete loss of SPOP. Similarly, the protein levels of Nup50, which only binds to SPOP from IVT-expressed proteins but neither co-immunoprecipitates with SPOP from HeLa cell lysates nor is degraded upon SPOP overexpression, also do not change upon SPOP knockdown. Consistent with these results, the use of siSPOP #101 and #102 resulted in an increase in the amount of Nup153 as detected via immunofluorescence microscopy (Figure 3G, H).

### SPOP may regulate Mad1 through Nup153

The nucleoporin Nup153 plays many roles in cell biology. One such role involves regulating the localization and thus function of Mad1, a spindle assembly checkpoint protein (Lussi et al., 2010). Lussi et al. showed that loss of Nup153 via siRNAs resulted in a weakened localization of Mad1 at the nuclear envelope, a stronger Mad1 association with the mitotic spindle, and cytokinetic defects (Lussi et al., 2010). We sought to determine whether loss of Nup153 via SPOP overexpression could also recapitulate the Mad1 phenotypes observed due to loss of Nup153 via siRNA knockdown.

Expression of neither SPOP^WT^ nor SPOP^F102C^ had any significant effect on the amount of Mad1 localized to the nuclear envelope in interphase (Figure 4A, B). Given that the weakened expression of Nup153 upon SPOP overexpression was only modest (a loss of about 40% by immunoblotting or about 50% by immunofluorescence), it is possible that SPOP overexpression and the subsequent modest loss of Nup153 is not a severe enough loss to phenocopy the Mad1 phenotypes previously observed due to loss of Nup153 by siRNA knockdown.

**Figure 4.**
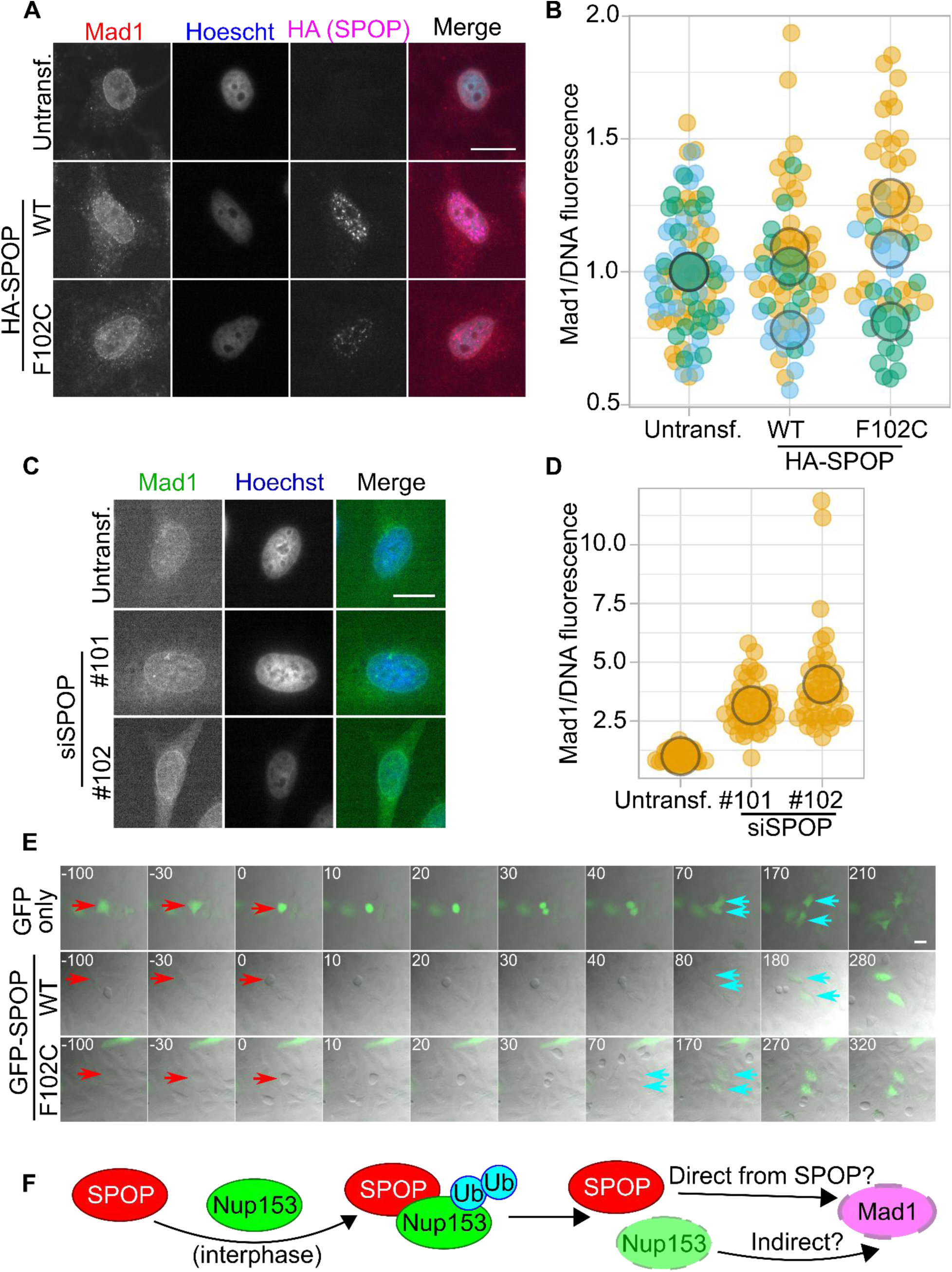
SPOP overexpression does not influence cell division. (A) HeLa cells transfected with HA-SPOP WT or F102C were imaged (z-stack) after staining with the indicated antibodies and (B) the ratio of nuclear enveloped-localized Mad1 to DNA fluorescence was quantified. (C) HeLa cells transfected with the indicated siRNAs against SPOP were imaged (z-stack) after staining with the indicated antibodies and (D) the ratio of nuclear enveloped-localized Mad1 to DNA fluorescence was quantified. (E) HeLa cells transfected with either GFP only or GFP-SPOP WT or F102C were visualized dividing after thymidine arrest. The time to mitotic rounding is given in minutes in the upper left corner of each frame. Red arrows point to the cell that will divide, while blue arrows point to the resulting daughter cells. (F) A model for the SPOP-Nup153 interaction. For (B) and (D), a different color indicates a different biological replicate, individual measurements are shown as a small dot, and the average value of a biological replicate is presented as a large circle. The scale bar in all microscopy images is 20 μm. Untransf., untransfected cells.

Since the loss of Nup153 resulted in a weakened localization of Mad1 at the nuclear envelope (Lussi et al., 2010), we wondered whether an increase in the amount of Nup153 would result in a greater amount of Mad1 at the nuclear envelope. Since loss of SPOP results in an increase of Nup153 at the nuclear envelope, we knocked down SPOP using siRNAs and quantified the amount of Mad1 at the nuclear envelope (Figure 4C, D). Similar to the increase of Nup153 at the nuclear envelope upon SPOP depletion, we saw an increase in the amount of Mad1 at the nuclear envelope, too.

Finally, we evaluated whether SPOP overexpression would have an effect on the progression of cell division. We overexpressed GFP, GFP-SPOP^WT^, or GFP-SPOP^F102C^ and observed the division of HeLa cells via microscopy (Figure 4E). No defects in cell division or timing were observed, although we did observe that cells had undetectable GFP-SPOP expression levels entering into mitosis but regained strong GFP-expression about 2.5 hours after cytokinesis, similar to what has been demonstrated previously via immunoblotting with cells at various stages of the cell cycle (Zhang et al., 2017a).

## Discussion

The nuclear pore complex is a crucial structure to scaffold other proteins and regulate nuclear trafficking. Here, we demonstrate that the Cul3 substrate adaptor SPOP binds to Nup153 and targets Nup153 for degradation. We also show that SPOP may regulate Mad1 localization to the nuclear envelope, though whether this regulation is mediated via Nup153, a direct SPOP-Mad1 interaction, or indirectly through another substrate, is unknown (Figure 4F).

Most SPOP substrates are degraded through the ubiquitin proteasome system, but recent reports have demonstrated that, at least in *Saccharomyces cerevisiae*, the nuclear pore complex is degraded through autophagy via recognition of Nup159 on the cytoplasmic side of the NPC (Lee et al., 2020; Tomioka et al., 2020). It is unclear how the protein stability of the nuclear pore complex is regulated in vertebrates. Here, we show that SPOP can degrade Nup153 but does not rule out autophagy as a means of degradation of the nuclear pore complex. Understanding the relationship between nutrient flux (for example, low nutrient conditions to stimulate autophagy) and SPOP-mediated degradation of Nup153 remains to be determined, as SPOP regulation can change under autophagy-inducing conditions. Indeed, SPOP mutants that no longer degrade BRD4 result in AKT and mTORC1 activation in prostate cancer cells and cell proliferation (Zhang et al., 2017b); perhaps the nuclear pore complex is degraded “in bulk” via autophagy during nutrient-poor conditions but Nup153 levels are “fine-tuned” via SPOP during nutrient-rich, proliferative conditions.

One possible piece of evidence to support this idea comes from the half-life of the NPC components: whereas NPC proteins in the central channel generally have a long half-life (approximately 200-700 hours in some cell types), Nup153 has a shorter half-life on the scale of about 50 hours (Mathieson et al., 2018). The observation that some NPC components are long-lived whereas Nup153 is a more dynamic and readily-turned over NPC component is also generally true in *C. elegans* models of aging (D’Angelo et al., 2009). Of the NPC protein components, Nup153 and Nup50 are the most dynamic, only transiently interacting with the NPC (Rabut et al., 2004) and perhaps are accessible to be regulated by SPOP. Thus, SPOP-specific mediated regulation of one component of the NPC, Nup153, may be one component of Nup153’s relatively rapid protein turnover. We note that while proteasomes have been shown to localize at the nuclear basket (where Nup153 resides) in yeasts and *C. reinhardtii* (Albert et al., 2017) and some mammalian NPC components are ubiquitylated (Chakraborty et al., 2008), we believe the SPOP-Nup153 interaction is the first identification of an E3 ubiquitin ligase for the nuclear pore complex in mammals. Generally, few E3 ubiquitin ligases have been recognized that target the nuclear pore complex: one example would be a Cdc53-Skp1-Grr1 complex (homologous to mammalian Cul1-Skp1-F box complex) that monoubiquitylates Nup159 in *S. cerevisiae* to control nuclear migration during mitosis (Hayakawa et al., 2012).

Indeed, how SPOP expression and activity can change in different cellular conditions, such as nutrient flux, is an important question for SPOP regulation. For example, in kidney cancers, SPOP increases during hypoxia and relocalizes into the cytoplasm where it gains a new suite of non-nuclear substrates (Li et al., 2014). During DNA damage, ATM kinase phosphorylates SPOP in the MATH domain and increases the affinity of SPOP for the substrates 53BP1 (Wang et al., 2021) and HIPK2 (Jin et al., 2021) (of note, these two SPOP substrates are ubiquitylated but not degraded). In proliferative, cycling cells, Aurora A kinase also phosphorylates SPOP in the MATH domain (Nikhil et al., 2020). While this phosphorylation was suggested to promote SPOP degradation (Nikhil et al., 2020), whether or not phosphorylation of SPOP in the MATH domain also allows SPOP to interact with a new suite of substrates is unclear. At least for one cell cycle-related kinase, Cdk4-mediated phosphorylation of SPOP resulted in association of SPOP with 14-3-3γ, an event that protected SPOP from Cdh1/Fzr1 binding and resulting degradation (Zhang et al., 2017a). Phenotypes related to SPOP-mediated degradation of Caprin1 were only evident when cells underwent stress and form stress granules (Shi et al., 2019). Finally, SPOP-mediated degradation of Cyclin E1 was only observed in some prostate and bladder cancer cells and not in the other tested cell types (Ju et al., 2018). Given the changing roles of SPOP under varied cellular conditions and cell type, there is a possibility that the effect of SPOP on Nup153 degradation may be stronger than the modest effect we see here or previously reported under other cellular conditions (Lan et al., 2019).

## Supporting information

STable1_Reagents

STable2_RawData

STable3_MS

Movie2_SPOPwt

Movie1_GFPonly

## Acknowledgements

We thank the members of the Torres lab for their assitance.

## Figures and Supplementary Materials

This manuscript contains:

4 main figures and legends; 3 supplemental figures and legends; attached are three supplemental files (.XLSX) and three movie files (.AVI).

SFile1 describes the plasmids, oligonucleotides, and antibodies (both for immunoblotting and immunofluorescence microscopy) used in this study.

SFile2 contains the raw data (numerical values) for all quantified values regarding fluorescence intensity, qPCR, and immunoblotting.

SFile3 contains the mass spectrometry data for pGLAP2-SPOP^WT^ in HEK293T cells.

Movie1, Movie2, and Movie3 contain the respective movie files of the GFP-only, GFP-SPOP^WT^, and GFP-SPOP^F102C^ cells dividing presented in Figure 4.

## References in Supplementary Files

Other plasmids were obtained from PMID: 26929214 (Cheung et al., 2016), PMID: 11448991 (Daigle et al., 2001), PMID: 15469984 (Hansen et al., 2004), and PMID: 25839665 (Xia et al., 2015).

## Figures and figure legends

**Supplementary Figure 1.**
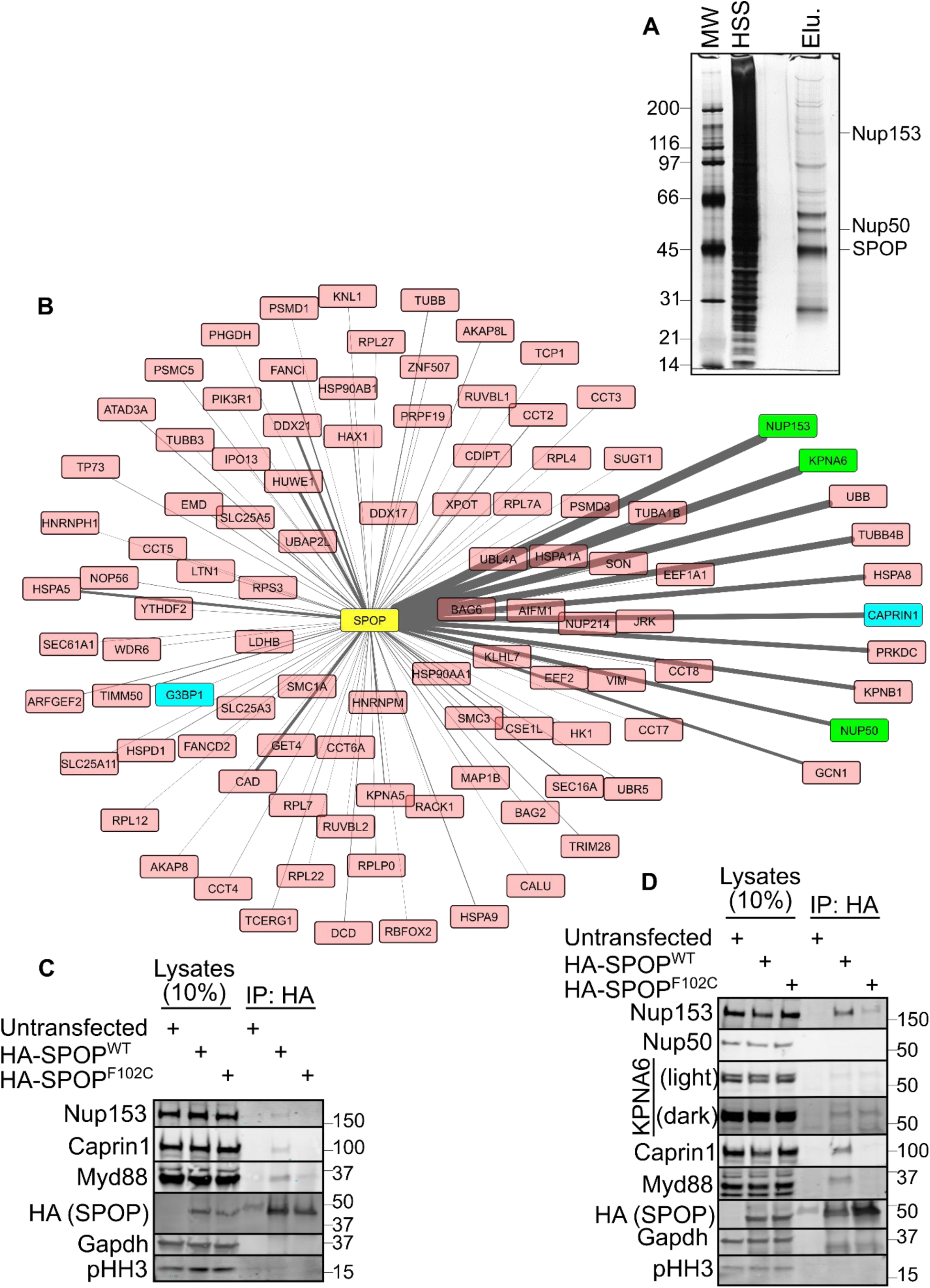
Identification and validation of SPOP-Nup153 binding interaction. (A) Coomassie-stained gel from tandem affinity purification of pgLAP2-SPOP from HEK293 cells. MW, molecular weight marker, given to the left in kDa. HSS, high spin supernatant. Elu., elution. Proteins of interest are noted on the right. (B) Cytoscape analysis of proteins detected by mass spectrometry. The thickness of the line is proportional to the peptide count. Known SPOP interactors are in blue, and SPOP interactors focused on in this study are in green. The ten most abundant hits are on the right. (C) Lysates from HeLa cells or (D) from HEK293T cells transfected with the indicated HA-SPOP plasmids were subjected to HA-immunoprecipitation and the resulting blots were probed with the indicated proteins. In (D), light and dark refer to low and high intensity scans.

**Supplementary Figure 2.**
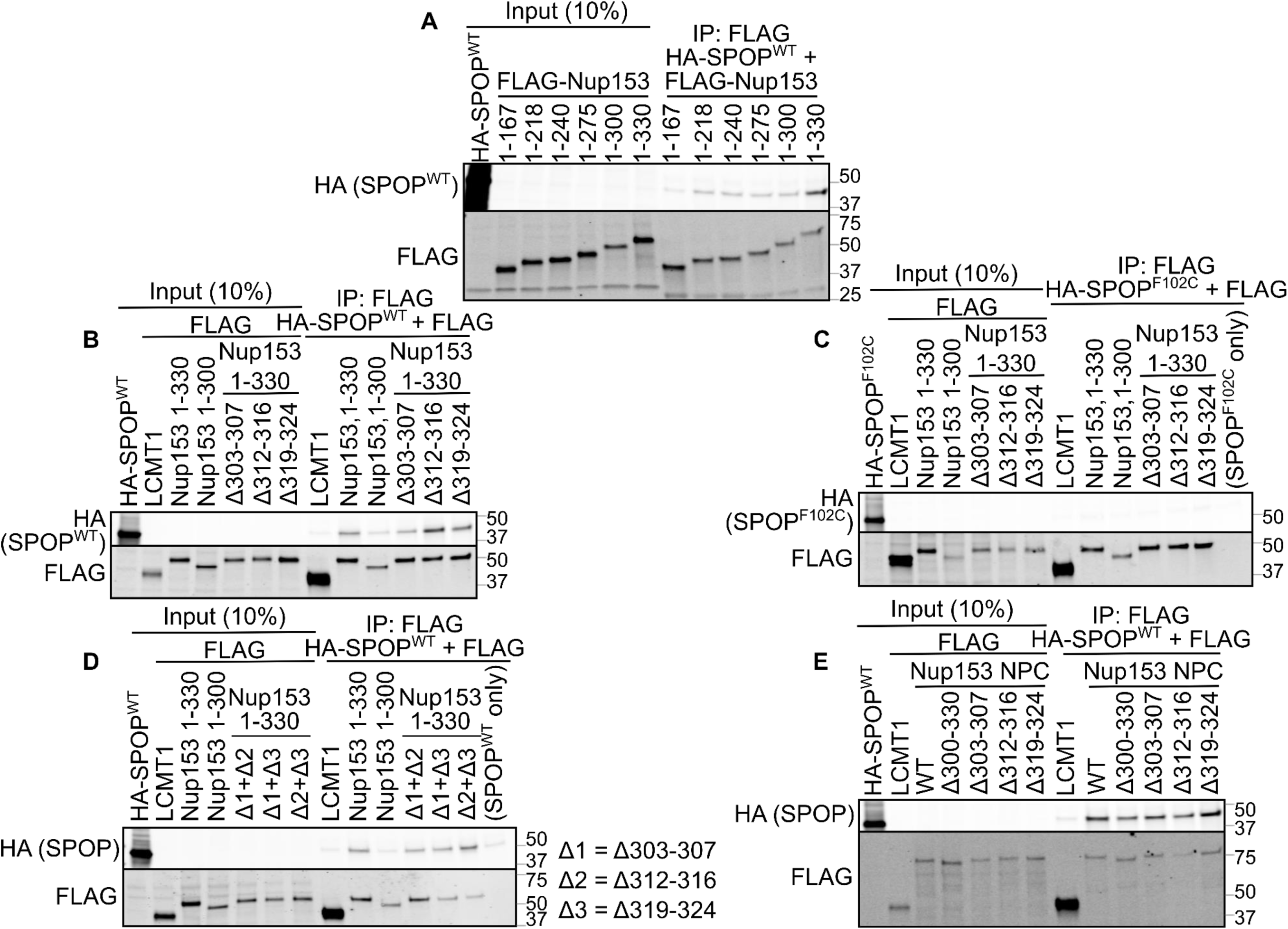
SPOP binds to Nup153 1-330 but not Nup153 1-300. (A) IVT-expressed HA-SPOP^WT^ and increasing N-terminal segments of FLAG-Nup153 were subjected to FLAG-immunoprecipitation. (B) IVT-expressed HA-SPOP^WT^ and (C) binding mutant HA-SPOP^F102C^ and FLAG-Nup153 1-330 with single deletions in the 300-330 region were subjected to FLAG-immunoprecipitation. (D) IVT-expressed HA-SPOP^WT^ and FLAG-Nup153 1-330 with double deletions in the 300-330 region were subjected to FLAG-immunoprecipitation. The double deletions in Nup153 are noted to the right of the immunoblot. (E) IVT-expressed HA-SPOP^WT^ and FLAG-Nup153 NPC with the indicated deletions were subjected to FLAG-immunoprecipitation. For all blots, FLAG-LCMT1 is used as a negative control and SPOP only refers to HA-SPOP and the FLAG beads (no FLAG-tagged protein).

**Supplementary Figure 3.**
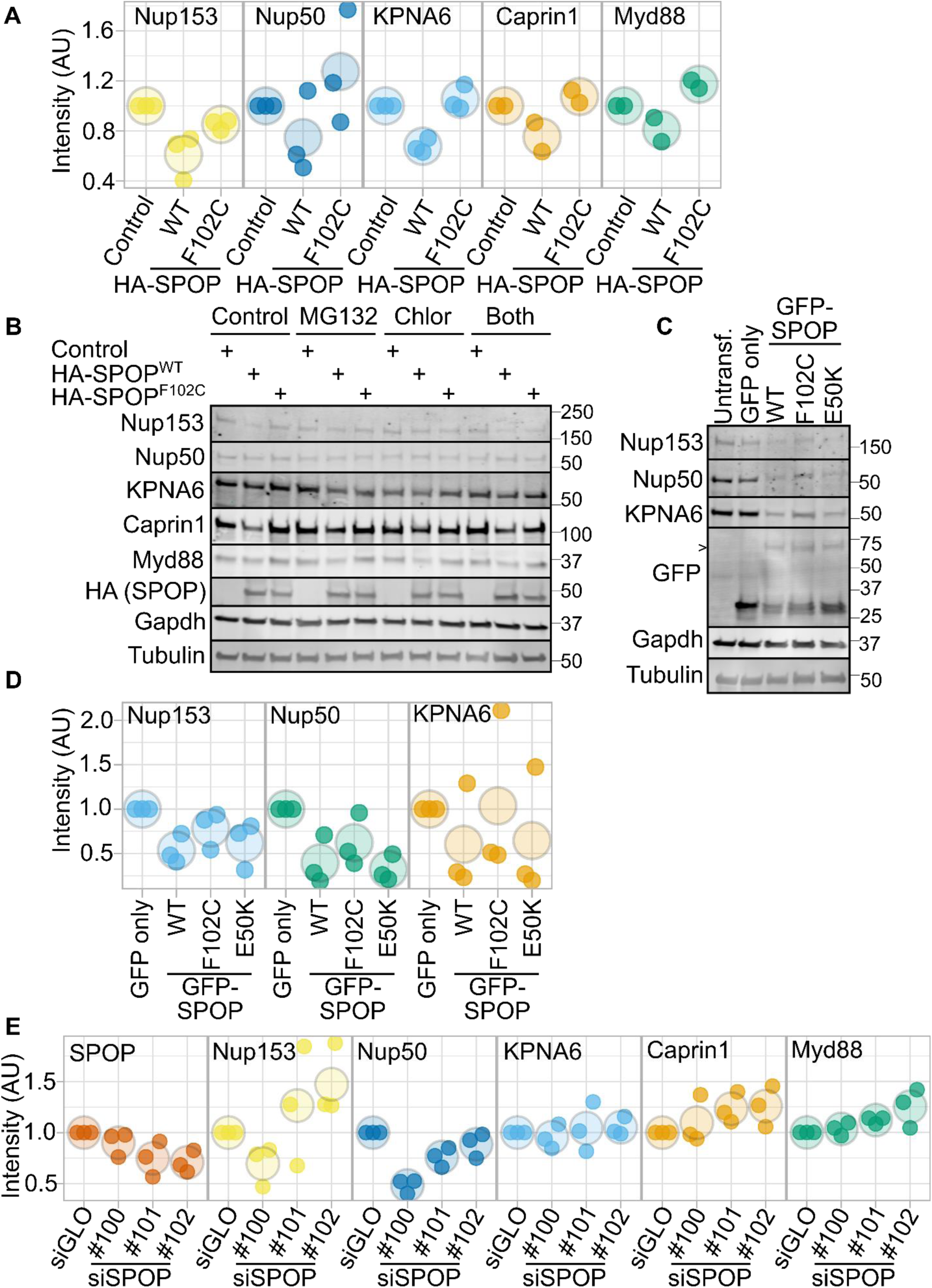
SPOP regulates Nup153 protein levels. (A) Quantification of the immunoblots (SPOP overexpression) shown in Figure 3A. (B) HeLa cells were treated with 0.2% DMSO by volume, 20 μM MG132, 50 mM chloroquine (Chlor), or both for 5 hours and the resulting lysates were probed with the indicated antibodies. (C) HeLa cells were left untransfected (untransf.), transfected with GFP only, or GFP-SPOP WT, F102C, or E50K and (D) the protein levels from the resulting immunoblots were quantified. The chevron arrow refers to GFP-tagged SPOP. (E) Quantification of the immunoblots (siRNAs against SPOP) shown in Figure 3F. For immunoblot quantification, all immunoblots were normalized to Gapdh and to the first condition in the figure, and a different color indicates a different biological replicate, individual measurements are shown as a small dot, and the average value of a biological replicate is presented as a large circle.

